# Somatic Structural Variant Discovery from Tumor Specific Strings Computed Against a Draft Composite Germline Reference

**DOI:** 10.64898/2026.11.06.764802

**Authors:** Davide Cesare Monti, Simone Ciccolella, Davide Cozzi, Gianluca Della Vedova, Paola Bonizzoni, Luca Denti, Fereydoun Hormozdiari

## Abstract

Somatic structural variants (SVs) are among the most clinically relevant genomic alterations in cancer. However, their accurate detection remains challenging due to complex genomic rearrangements, repetitive genomic regions, difficulties in distinguishing somatic variants from germline variation and sequencing artifacts, and limited sequencing support for low-frequency events. Existing mapping-based somatic SV callers are susceptible to alignment ambiguities in repetitive regions and near complex SV breakpoints, while approaches relying on comparisons of independently generated tumor and normal variant callsets may propagate errors from both callsets. Building on the concept of *specific strings*, previously applied to germline SV discovery, we introduce `VESPA-SS`, a novel approach for detecting somatic SVs from paired tumor–normal long-read whole-genome sequencing data. `VESPA-SS` directly extracts somatic SV signals using specific strings computed from tumor reads against a *Draft Composite Germline Reference*, without requiring initial read mapping or a high-quality germline assembly. These signals are subsequently clustered and refined through consensus sequence generation and alignment-based procedures to identify somatic SVs. Evaluation on real PacBio HiFi datasets demonstrates that `VESPA-SS` outperforms state-of-the-art somatic SV callers, achieving F1 scores exceeding 0.92 and 0.90 on the well-characterized HG008 and COLO829 cancer cell lines, respectively, compared with the highest F1 scores of 0.86 and 0.85 achieved by the evaluated methods. These results demonstrate the effectiveness of specific string based signal extraction for accurate somatic SV discovery and establish a new computational paradigm for identifying complex genomic alterations in cancer. `VESPA-SS` is freely available at https://github.com/dcmonti/VESPA.

## 1 Introduction

Somatic structural variants (SVs) are a defining feature of cancer genomes. Mutational processes in tumor are guided by deletions removing tumor suppressors, duplications and amplifications raising the dosage of oncogenes, and balanced and unbalanced rearrangements creating gene fusions and reshaping chromosomes, often through complex events such as chromothripsis that join many breakpoints at once [17]. Accurate detection of these variants is critical for understanding the mutational processes that drive the tumor evolution and for guiding clinical decisions that rely on the identification of specific genomic rearrangements. Short reads systematically miss entire classes of somatic SVs, such as large-scale genomic rearrangements, events in repetitive regions of the genome, and variants with clustered breakpoints [11, 6]. On the contrary, third-generation sequencing technologies, such as Oxford Nanopore and PacBio, help overcome these limitations by producing long reads that span structural variant breakpoints and can be phased across extended genomic regions, providing direct evidence for large and complex structural rearrangements [11, 6]. As a result, accurate long reads, such as PacBio HiFi [31], can, in principle, enable the comprehensive characterization of the somatic SV landscape of a tumor at base-pair resolution.

Existing approaches typically call somatic SVs by aligning tumor and matched-normal long-read sequencing reads to a reference genome and detecting structural variants from the resulting read alignments. `Severus` builds a phased breakpoint graph from the alignments of both samples [11], `SAVANA` classifies breakpoint clusters with a model trained on long-read cancer data [6], `nanomonsv` compares tumor and normal breakpoint evidence [25], and germline callers, such as `Sniffles`, are applied in mosaic mode with normal calls subtracted from tumor ones [26]. All existing methods compare tumor and matched-normal read alignments to a generic reference genome rather than to the patient’s own germline genome. As a result, germline variants absent from the reference appear as candidate SVs in the tumor and are recognized as germline, and thus excluded from the somatic call set, only when sufficient evidence is also observed in the matched-normal sample. This paradigm implicitly assumes that tumor and normal reads covering the same locus are aligned similarly and that sufficient coverage is available in both samples to observe and compare the corresponding alleles reliably. However, this assumption often breaks down in complex genomic regions, such as tandem repeats and segmental duplications, where alignments may differ between the two samples despite identical underlying sequences. Combined with the low allele fraction of somatic events, which is diluted by tumor purity and subclonality, this makes false positive calls a major limitation of current methods. Consequently, the somatic SV call sets produced by different callers often show limited concordance [14].

A natural solution is to replace the reference genome with a patient-specific germline genome, enabling somatic variants to be identified directly through tumor-germline comparison. Recent studies replace the reference genome with a diploid assembly of the matched-normal sample and report substantial gains in both precision and recall of somatic SVs, especially in complex regions of the genome [32, 19, 1]. Others use a self-assembly of the normal to filter calls whose supporting reads align without the SV against it [23], or co-assemble the tumor and the normal and call long-range SVs from the topology of the joint assembly graph [14]. However, their performance is generally influenced by the assembly being a faithful, high-quality, and nearly complete representation of the germline genome. Near-telomere-to-telomere diploid assemblies currently are obtained from high-coverage, multi-platform sequencing, typically combining PacBio HiFi with ultra-long Oxford Nanopore reads [27, 19], a sequencing effort far beyond that of a typical tumor–normal study. At moderate sequencing coverage, germline assemblies are often fragmented and incomplete, making regions absent from the assembly potential sources of false somatic calls. Furthermore, using a donor-specific assembly as the reference requires aligning both tumor and normal samples to each haplotype, calling variants separately, and reconciling the calls back to a common coordinate system [32].

In this work, we explore a different direction based on the notion of *sample-specific strings* (SFSs) [12]. By directly capturing sequence differences between samples, specific strings provide a framework for somatic SV discovery that can, in principle, circumvent many of the limitations of current approaches. Given a target and a reference set of strings, a sample-specific string (or simply specific string) is a shortest string that occurs in the target but not in the reference [12]. A specific string, therefore, marks directly at the sequence level and without alignment where one sample differs from another. We previously used SFS to compare samples [12], to call germline SVs in hard-to-call regions of the genome [4], and to augment pangenome graphs from unassembled long reads [3]. The somatic setting is a natural fit for this notion: the sequence introduced by a somatic event is, by definition, present in the tumor genome (and reads) and absent from the patient’s germline genome (and reads). This observation has an important practical consequence. Identifying a comprehensive set of tumor specific strings that captures somatic evidence does not require a high-quality assembly of the patient’s germline genome. Instead, it is sufficient that the patient’s germline sequence is represented, even in fragmented form, within the collection of sequences against which the tumor reads are compared.

We therefore introduce the concept of *Draft Composite Germline Reference*, which can be intuitively viewed as a reasonably complete representation of the patient’s germline sequence. This reference combines a moderate-coverage *de novo* assembly of the normal sample, the reference genome, and the normal reads containing sequence not represented in either of the first two components. This combination provides a near-complete representation of the germline genome without requiring a costly high-quality assembly. The underlying rationale is that any germline sequence omitted from this representation will appear as a spurious tumor specific string, thus generating biologically uninformative evidence for somatic SV discovery. This framework enables the alignment-free extraction of somatic evidence directly from paired tumor–normal long-read whole-genome sequencing data, addressing many of the limitations of existing approaches. We build upon and substantially extend our previous framework for germline SV discovery [4] through four key innovations: (i) a somatic-aware error-correction strategy that preserves low-frequency somatic signals, (ii) a haplotype-informed non-transitive clustering algorithm for specific strings, (iii) support for the detection of large somatic rearrangements, and (iv) probabilistic filtering procedures that improve SV calling precision by identifying candidate somatic events that are likely germline.

Building on these innovations, we present `VESPA-SS` (Variant Extraction from SPecific strings with draft Assemblies - Somatic Structural variants), a method for the discovery of somatic structural variants from PacBio HiFi sequencing of paired tumor and matched-normal samples that incorporates the above innovations. We evaluated `VESPA-SS` on two cancer cell lines sequenced with PacBio HiFi. Specific strings exclude most germline variation, enabling `VESPA-SS` to call somatic events with low Variant Allele Frequency without sacrificing its precision. This preserves subclonal events and gives `VESPA-SS` the highest *F*_1_ on both datasets. Moreover, `VESPA-SS` does not require a high-quality germline assembly: a Draft Composite Germline Reference built from HiFi reads of the normal at moderate coverage yields the same true positives as a curated, near telomere-to-telomere germline assembly of the same individual, with only one additional false positive.

## 2 Methods

In this section, we describe `VESPA-SS`, our novel framework for somatic SV discovery from paired tumor–normal long-read whole-genome sequencing data based on specific strings. An overview of the proposed approach is illustrated in Figure 1. `VESPA-SS` consists of four main steps. First, tumor and matched-normal long reads are error-corrected using a strategy designed to remove sequencing errors while preserving low-frequency somatic variants (Section 2.1). Second, a Draft Composite Germline Reference is constructed by combining a de novo assembly of the matched-normal sample, the reference genome, and normal reads containing sequences not represented in either. Tumor specific strings are then computed against this Draft Composite Germline Reference to identify sequence differences potentially associated with somatic events (Section 2.2). Third, the specific strings are localized to the reference genome using the alignments of their supporting reads, then clustered and analyzed to reconstruct candidate somatic SVs, including large and complex genomic rearrangements (Section 2.3). Finally, candidate SVs are evaluated against the matched-normal reads using a series of filters designed to distinguish genuine somatic events from germline variation and assess the reliability of the supporting evidence (Section 2.4). We note that `VESPA-SS` exposes several user-defined parameters that control its execution. Unless otherwise specified, the methodology described in the following sections assumes these default parameter settings.

**Figure 1.**
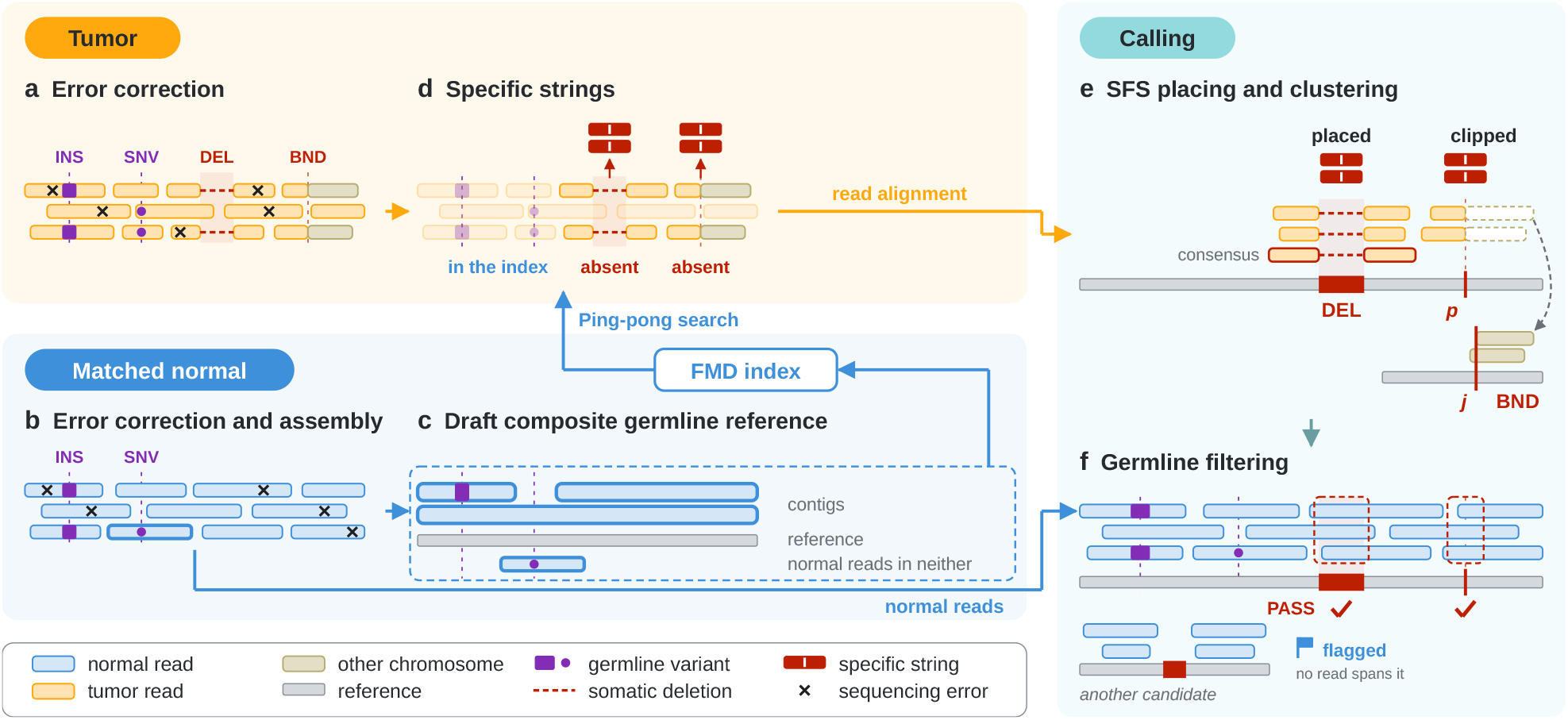
**a, b** Tumor and normal HiFi reads are error-corrected, removing sequencing errors and preserving germline and somatic events; the normal reads are also assembled. **c** The Draft Composite Germline Reference (normal contigs, reference genome, and the normal reads that neither contains) is indexed in an FMD-index. **d** The specific strings (SFS) of the tumor reads are computed against it: germline variants yield none, somatic events do. **e** SFS are placed on the reference through the alignment of their reads; placed SFS are clustered and called from a consensus, clipped SFS give a breakpoint *p* whose partner *j* comes from the supplementary alignment. **f** Candidates are compared with the normal reads spanning them: those spanned but not carried are marked `PASS`, those that no normal read spans are flagged.

### 2.1 Somatic variant preserving error correction

Specific strings capture every difference between the tumor reads and the Draft Composite Germline Reference, including sequencing errors. An error in a tumor read yields an SFS that supports no real variant between the tumor and the normal. Since errors occur independently in every read, such SFS are spread over the whole genome and far outnumber those induced by somatic events. Sequencing errors in the normal reads can affect the analysis in two ways. First a normal read carrying an error can enlarge the Draft Composite Germline Reference without adding any real germline sequence. Second, errors in normal reads spanning the breakpoints of a candidate SV may appear as spurious indels in their alignments, which can be incorrectly interpret as evidence that the event is also present in the normal sample. To minimize these effects, we error-correct reads from both tumor and matched-normal samples before computing specific strings.

A promising starting point is the alignment-free error-correction approach introduced in `hifiasm` [2] and later utilized in palss [3]. In `hifiasm`, overlapping reads are classified based on their consistency at informative heterozygous positions, allowing only reads inferred to originate from the same haplotype to contribute to error correction. Although this haplotype-aware strategy helps preserve germline heterozygous variation, the majority-based correction procedure may inadvertently eliminate low-frequency somatic variants supported by only a minority of reads, treating them as sequencing errors. To address this limitation, we modify the correction procedure to preserve bases supported by a sufficient number of reads, even when they do not represent the majority. Rather than applying a fixed support threshold, we determine the minimum required support using a negative binomial model of somatic read support and a user-specified tolerance for the loss of somatic events.

Let *C* be the expected coverage of the tumor sample and *ρ* its purity, i.e., the fraction of tumor cells in the sample. A somatic event carried by a fraction *f* of the tumor cells, at a locus of copy number two, is shown on average by 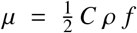reads, the factor one half because the event lies on one of the two haplotypes. Here *ρ* is a property of the sample, whereas *f* varies from event to event: we take *f* as the smallest fraction of tumor cells whose events we want to preserve, since events carried by more cells have more supporting reads and are lost less often. We then derive the required support *T* from µ and from the fraction φ of such events the user accepts to lose, instead of fixing it for each depth, purity, and subclonal fraction. The error correction can remove an event only if fewer than *T* reads support it, so the fraction of somatic events it loses is at most Pr (*X* < *T*), with *X* the number of reads supporting an event. Read depth varies along the genome and is commonly modeled as a negative binomial [18, 21]. Each read covering the event carries it independently with probability *ρf*/2, and such a binomial thinning of a negative binomial is again a negative binomial with the same over-dispersion. We therefore model *X* as a negative binomial with mean µ and variance µ α+µ^2^, where α is the over-dispersion of the read depth, and take *T* as the largest threshold whose loss is below φ:

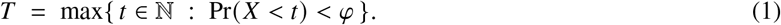

The over-dispersion α depends on the sequencing run; we measure it on the normal samples of this study.

### 2.2 Draft Composite Germline construction and specific strings computation

To represent the germline genome, we introduce the notion of *Draft Composite Germline Reference*, from now on simply *germline set*. Since any sequence present in the tumor reads but absent from the index yields an SFS, whether it represents a somatic event or not, the germline set must contain as much of the germline variation as possible. For this reason, the germline set is made of three components. The first is a de novo assembly of the matched normal sample, obtained with `hifiasm` [2]. However, an assembly obtained at moderate depth is typically not complete, and a tumor read crossing a region absent from the assembly would be reported as specific string, even though it does not support any somatic event. Therefore, the second component is the reference genome, which covers the regions where the reads of the normal are too sparse to be assembled. However, germline variants not represented in the assembly are also absent from the reference genome. To cover these variants, the third component consists of the reads of the normal sample that contain at least one specific string with respect to the assembly and the reference genome. To compute specific strings, we use the Ping-Pong algorithm introduced in [12], which efficiently identifies such strings using an FMD-index [16]. The index of the germline set is therefore built in two steps. We first index the assembly and the reference genome, and compute the SFS of the corrected normal reads with respect to this index. The normal reads with at least one SFS are then added in their entirety to the index. The resulting index is used to compute specific strings from the tumor reads in the next step of the method. Additionally, to address homopolymer runs reported with an incorrect length, the dominant error mode of HiFi reads [31], we truncate every homopolymer run longer than a user-defined cap, both in the germline set before indexing and in the reads, as done by HiFi assemblers [22]. Specific strings are computed on the compressed sequences and mapped back to the original reads.

### 2.3 Candidate calling from specific strings

A specific string marks the part of a tumor read that differs from the germline set, but it gives neither the position of the event on the reference genome nor its breakpoints, type, or length. Calling SVs from SFS therefore requires mapping each SFS on the reference genome, grouping the SFS supporting the same event, and reconstructing the event from each group. An SFS is mapped through the alignment of its read, by projecting its start and end from the read onto the reference [4]. We denote an SFS as *placed* if it lies within the aligned part of its read, so that both ends can be projected, and *clipped* if it reaches into a soft-clipped part of the read. In SVDSS [4], SVs are called exclusively from placed SFS, clustered using a greedy approach, where an SFS is added to the current cluster if it overlaps the region spanned by the cluster, which is progressively expanded as new SFSs are added. Although computationally efficient, this greedy approach can inadvertently merge nearby distinct events when a long SFS spans both events, causing all SFSs supporting either event to be grouped together and potentially resulting in spurious or missed SV calls. Furthermore, clipped SFS are discarded in SVDSS [4], despite potentially providing evidence of large genomic rearrangements, which are particularly prevalent in the somatic setting [17]. To address these limitations, we employ two complementary strategies for SV calling from SFSs. Placed SFS are grouped by a novel clustering algorithm (Section 2.3.a), while clipped SFS are clustered by the junction they support (Section 2.3.b). In both cases, `VESPA-SS` retains only clusters of specific strings supported by at least *s* reads.

#### 2.3.a Clustering and consensus of placed specific strings

We cluster placed specific strings using a novel algorithm based on hierarchical agglomerative clustering (HAC), in which two SFSs are assigned to the same cluster only if they are sufficiently close to each other on the reference genome. This prevents long SFSs spanning multiple nearby events from merging otherwise distinct clusters.

We define the distance between two placed SFSs, *A* and *B*, based on the overlap of their reference intervals, [*a*_*s*_, *a*_*e*_] and [*b*_*s*_, *b*_*e*_], respectively:

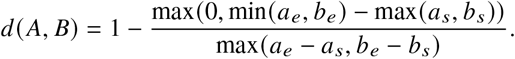

This distance ranges from 0 to 1 and measures the extent of overlap relative to the longer interval. Normalizing by the longer interval ensures that SFSs with substantially different lengths are not considered close, preventing long SFSs from connecting distinct events.

Initially, each SFS forms its own cluster. At each subsequent iteration, the two closest clusters are merged, with the distance between clusters defined using complete linkage, i.e., the maximum pairwise distance between SFSs belonging to the two clusters. Since complete linkage produces a non-decreasing sequence of merge distances, clustering terminates when the minimum distance between any two remaining clusters reaches 1, indicating that any further merge would combine non-overlapping SFSs. We then identify the largest increase in the recorded merge distances and select the clustering obtained immediately before this increase as the final clustering. A cluster, however, may contain the reads of two alleles of the same locus, e.g., a heterozygous event. The two alleles occupy the same reference interval, so the clustering cannot separate them, and an event called from the whole cluster would wrongly merge the two. We therefore split the reads of each cluster by their haplotype tag (when available), which is assigned to the whole read and thus provides longer-range information than the cluster region itself. Untagged reads join the subcluster of closest length, and when no read is tagged, the cluster is split into up to two subclusters by length alone. For each subcluster, we reconstruct the underlying putative SV event by generating a consensus sequence from its supporting reads. The reads are first trimmed to a common interval surrounding the SFSs in the cluster and aligned into a partial order graph with abPOA [9]. Because missing or incorrect haplotype tags may cause a subcluster to contain reads supporting two distinct alleles of similar size, we allow abPOA to generate up to two consensus sequences. Specifically, reads are partitioned into two groups based on sequence differences represented in the alignment graph, and a separate consensus is constructed from each group to preserve allele-specific variants. Each resulting consensus is then aligned to the corresponding reference interval, defined by the boundaries of the cluster, using ksw2 [15]. Finally, SVs are identified from the gaps in these alignments.

#### 2.3.b Junction reconstruction from clipped SFS

A large genomic rearrangement can bring together two regions of the reference genome at positions known as *breakpoints*, forming a novel *junction* absent from the reference. Reads spanning such junctions typically exhibit split alignments, often with *soft-clipped* sequences at the breakpoints. A clipped SFS covering a junction therefore has one end in the clipped portion of the read; in the following we call it simply a *clip*. We note that SFS lying entirely in the soft clipped portions of the reads are discarded. A clip is a *left clip* when it starts in the clipped portion of a read and ends in its aligned part, and a *right clip* in the symmetric case. The breakpoint *p* of a clip, on chromosome *c*, is the reference position where the alignment of the read starts, for a left clip, or ends, for a right clip. Computing clips from specific strings and not directly from read alignments allows us to consider only a subset of the clipped reads, and specifically those that are clipped because of a somatic event. In fact, a read clipped due to a germline difference from the reference spells a sequence that the germline set contains and yields no specific strings.

A clip, however, locates only one of the two breakpoints of a junction and pairing clips by distance would infer only events with close breakpoints. Instead, we leverage the alignment of the clipped part itself, which is typically reported as a *supplementary alignment*. Among the supplementary alignments of the read, we call *partner* of the clip the one closest to the clip on the read. The other breakpoint of the junction is the position *j*, on chromosome *c*^′^, where the alignment of the partner starts or ends, depending on the type of the clip. The junction is thus described by the *clip breakpoint* (*c, p*), the *partner breakpoint* (*c*^′^, *j*), and its *orientation*: *direct* when the partner is aligned on the same strand as the primary alignment, *opposite* otherwise. Clips are then clustered by position: a clip joins a cluster when it lies within a fixed distance of the clip that opened it, and opens a new cluster otherwise. Left and right clips are clustered separately, because a left and a right clip at the same position never belong to the same junction. A cluster may collect clips of nearby junctions, or clips whose supplementary alignment is misplaced. We therefore split it into subclusters of clips whose partners agree, and take the number of clips of a subcluster as its weight. Finally, structural variants are called from each subcluster, inferring their type and length from the relative position of the two breakpoints of its junction, from its orientation, and from the read bases between the clip and its partner (as illustrated in Figure 2). We note that the same event may be represented by two different clip-partner pairs, depending on how the aligner maps the reads. In such cases, we report the event only once and sum the support contributed by both pairs.

**Figure 2.**
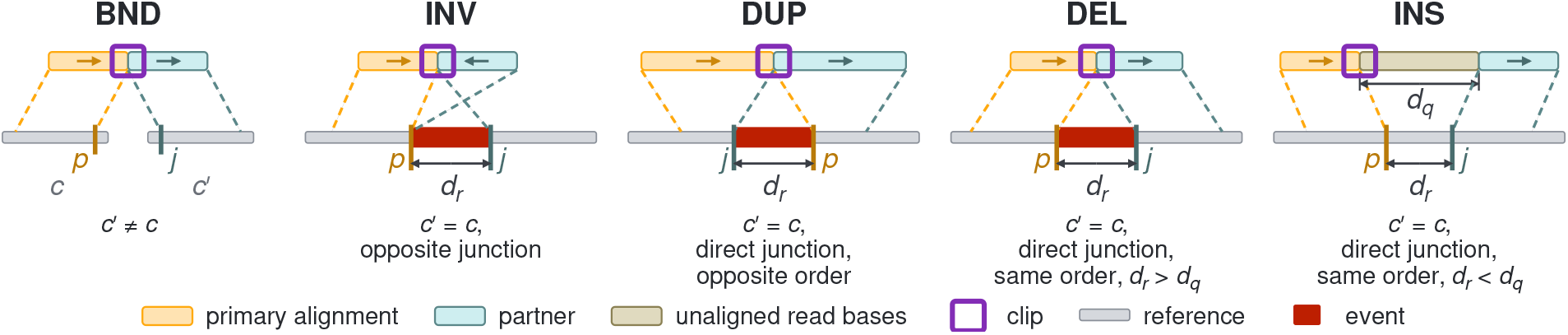
From the junction of a subcluster to the event, shown on one read crossing the junction. Dashed lines link each aligned part of the read to the interval where it aligns, and arrows give its strand relative to the primary alignment. *d*_q_ is averaged over the reads supporting the junction, and the order is *same* when *p* and *j* lie on the reference in the order of the clip and the partner on the read. Under each panel, the condition that yields the event; INV, DUP and DEL have length *d*_r_, and INS has length *d*_q_ − *d*_r_.

### 2.4 Assessing the somatic origin of candidates

Specific strings mark where the tumor reads differ from the germline genome of the patient, but each SV is called against the reference genome. A small difference between the two samples, such as a few extra units of a tandem repeat, can thus produce a call that is mostly germline. Germline sequence absent from the normal reads, e.g., for lack of coverage, also yields specific strings and thus a candidate variant. Each candidate SV is therefore tested against the normal reads that span its breakpoints, first for the presence of the event and then for whether its absence is informative.

Three filters test the presence, comparing the call with the indels of the normal reads near its breakpoints under a length criterion that is relaxed at each step. The same-allele filter discards the call when a normal read carries the same allele. The germline filter flags it when more than one normal read carries an indel of similar length. The polynormal filter flags it when many normal reads carry an indel of the same type, whatever its length. The repeatdepth filter flags the call when the normal depth at its breakpoints is far above that of the sample, a sign of reads mapped from copies of a repeat missing from the reference [29].

#### Power of the normal at the locus

Conversely, the normal reads at a locus may be insufficient to show a germline event. The lowpower filter estimates the probability of such a missed evidence, based on the number of normal reads that could carry the event and the likelihood that each of those reads would show it.

A read that carries the event does not always show it in its alignment, e.g., when the event is collapsed in a tandem repeat or the read is mismapped. We therefore let *q*_i_ be the probability that normal read *i* shows a germline event that it carries, and *q* the mean of the *q*_i_. If *D* normal reads carry the event, none of them shows it with probability at most (1 − *q*) ^D^, under the assumption of independence. Since a heterozygous germline event is carried only by the normal reads of its haplotype, *D* has to count only the normal reads matching the haplotype of the call. This is possible because tumor and normal reads are both tagged from the phased variants of the normal, so that a tag names the same haplotype in both samples. The haplotype of the call is thus the one carrying most of its tagged supporting reads, and *D* is the normal depth on that haplotype. Since part of the normal reads are untagged, this depth is estimated for each haplotype *h* as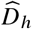, adding to its tagged normal reads spanning the locus a share of the untagged ones, proportional to the tagged counts with Laplace’s rule of succession [13, 10]. Combining the two quantities, we estimate an upper bound on the probability *P*_miss_ of missing the event had it been germline as 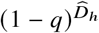 on the haplotype *h* of the call. When the haplotype of the call is undetermined, *P*_miss_ is estimated by averaging over the two haplotypes. The filter flags the call when *P*_miss_ exceeds a user-defined threshold.

## 3 Results

We evaluated `VESPA-SS` on two publicly available paired tumor–normal long-read whole-genome sequencing datasets generated using the PacBio HiFi Revio system, each accompanied by published somatic SV call sets used here as ground truths. The `HG008` dataset is a cell line of a pancreatic ductal adenocarcinoma, with pancreatic tissue of the same patient as normal, sequenced at 79× (tumor) and 35× (normal) [20]. The ground truth is based on the GIAB somatic SV draft benchmark v0.5 [30], which includes 145 clonal SVs and 173 SVs when both clonal and subclonal events are considered, within benchmark regions covering respectively 95.5 and 95.6 % of the genome outside the gaps. COLO829 is a cell line of a metastatic melanoma, with a lymphoblastoid cell line of the same donor as normal, sequenced at 63× and 66× ; its ground truth comprises 62 events across the entire genome [8]. The sequencing datasets are aligned to the `GRCh38` reference genome [24] with minimap2 [15] using its HiFi preset. As recommended in [23], the COLO829 dataset is also aligned setting -s50, which lowers the minimum alignment score threshold and allows shorter supplementary alignments to be reported. For `VESPA-SS`, homopolymer runs are truncated to two bases and SFS longer than 500 bp are discarded. The source code of `VESPA-SS`, along with scripts and instruction for reproducing the experimental evaluation, is available at https://github.com/dcmonti/VESPA.

### 3.1 VESPA-SS outperforms State-of-the-Art Somatic SV Callers

We first evaluated the accuracy of `VESPA-SS` in calling somatic SV from long-read sequencing datasets. To put our results in perspective, we compared `VESPA-SS` with four top-performing somatic SV callers: Severus [11], SAVANA [6], nanomonsv [25] and Sniffles2 [26]. We ran each state-of-the-art caller on the original, uncorrected reads, at its current release, with the command lines of the `HG008` Minda ensemble, from which the candidate SVs of the `HG008` ground truth were selected [30].

To assess the effect of the minimum support threshold for reporting a call on each caller’s accuracy, we performed a grid search over this parameter (*s* ∈ {3, 5, 8, 10, 15, 20, 25}). In addition to our own runs of these callers, we included, as external references, the call sets published from the same sequencing runs: on `HG008`, those of the Minda ensemble; on COLO829, the Severus call set of [11] and those of Qin et al. [23], including the call sets of minisv. colorSV [14] is not included, since it calls only long-range SVs. In our evaluation, we considered only calls marked `PASS`, and SVs shorter than 50 bp are removed from both the ground truth and the call sets. The accuracy of each caller was evaluated using Minda [11], which computes precision, recall, and F1 score through breakpoint-based matching between events in the ground truth and call set. We additionally benchmarked all callers using truvari [7]. As the two evaluation frameworks produced highly consistent results, we focus here on the Minda-based evaluation. As recommended by GIAB for the `HG008` ground truth, we restrict its evaluation to the benchmark regions (i.e., clonal and subclonal): calls falling outside them are not considered. Results of this evaluation are shown in Figure 3.

**Figure 3.**
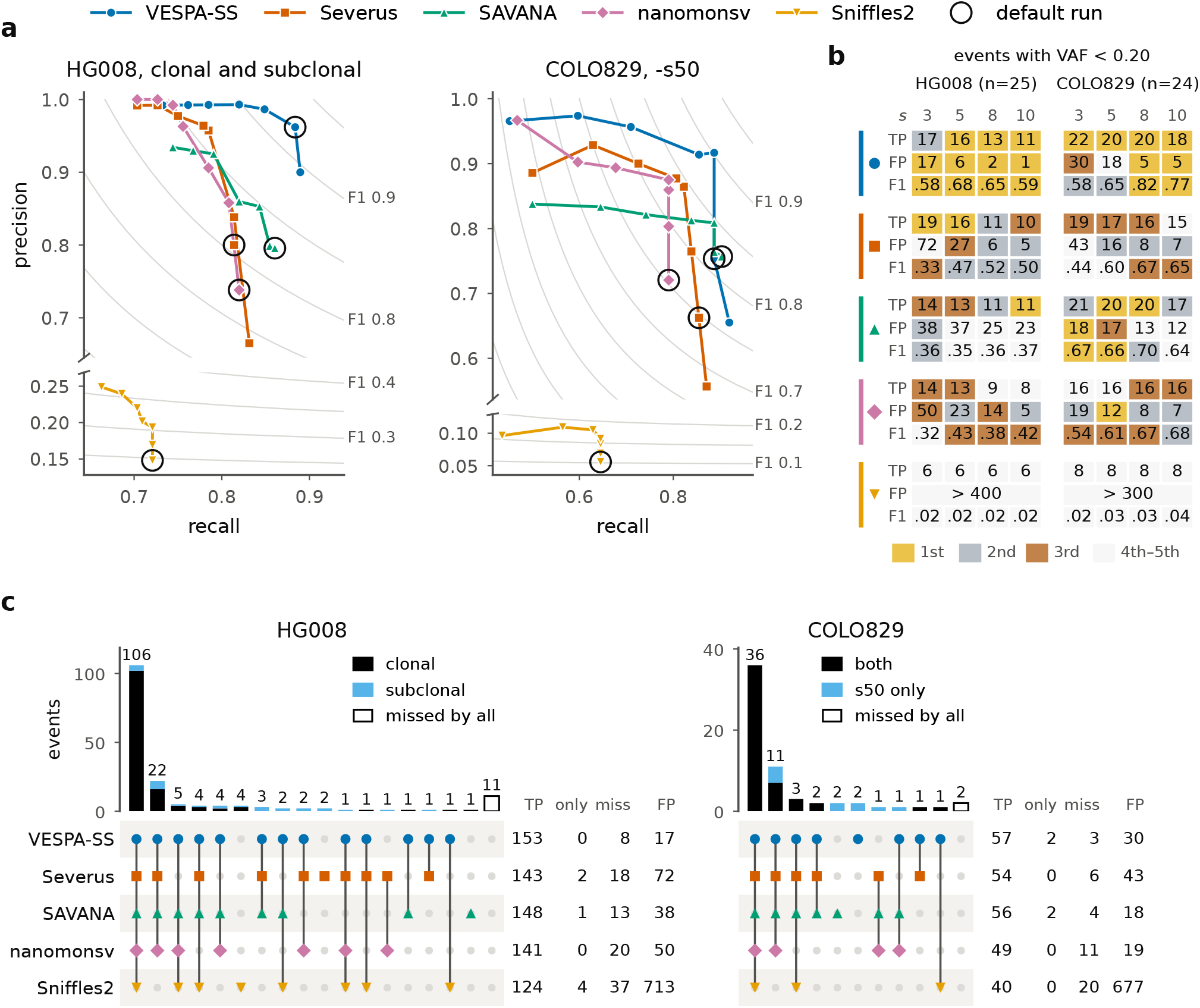
Comparison with the other callers, evaluated with Minda. (a) Precision and recall of every run of the grid; each line joins the runs of one caller from *s* = 3, at its highest recall, to *s* = 25. The default run of Severus lies off its line, since an explicit minimum support also disables its VAF filter. The precision axis is broken, with the same scale in both segments; grey lines are *F*_1_ isolines. (b) Events with a VAF below 0.20; callers are marked as in (a). TP, the events of the class found by the run; FP, the false positives of the whole call set; F1, 2 TP 2 TP FP FN, with FN the events of the class not found. Cells are colored by the position of the caller at that threshold (most TP, fewest FP, highest F1); tied callers share the better position. (c) Ground-truth events found by each caller at *s* = 3, on the data of (a). Each column is the set of events found by exactly the callers marked below it; on `COLO829`, s50 only marks the events that no caller finds with the default alignment (-s200). On the right: only, the events found by this caller alone; miss, the events found by another caller and not by this one; FP, the false positives of the run, which cannot be assigned to an intersection.

Overall, across all tested datasets and values of *s*, `VESPA-SS` consistently achieved the highest F1 score, with only few exceptions on the `COLO829` dataset. As expected, increasing the minimum support *s* required to report a call improved precision for all callers at the expense of their recall. When considering the best-performing configuration of each caller (i.e., the *s* value yielding the highest F1 score), on the `HG008` dataset, `VESPA-SS` with *s* = 5 achieves the highest F1 score (0.921) followed by Severus (*s* = 8) that exhibits an F1 score of 0.863, an increase of 5.9 percentage points. In comparison, the best published call set, SAVANA from the Minda ensemble, achieves an F1 score of 0.834. In particular, `VESPA-SS` achieves the highest recall, with a precision second only to that of nanomonsv (0.962 against 0.992). As expected, the performance of all callers improves when restricting the evaluation to the less challenging clonal regions. In this setting, the advantage of `VESPA-SS` is reduced but remains evident. `VESPA-SS` (*s* = 8) achieves an F1 score of 0.935, compared to 0.910 for both Severus and nanomonsv (*s* = 20), which each find 121 events with no false positives. A similar trend is observed on the `COLO829` dataset, where `VESPA-SS` (*s* = 8) achieves the highest precision and, tied with SAVANA (*s* = 8), the highest recall, resulting in an F1 score of 0.902, 5.5 percentage points above SAVANA and 5.2 above the best published call set, SAVANA filtered with minisv (0.850). With the alignments generated with the default minimap2 parameters, the F1 score of all callers but Sniffles2 decreases by 0.06 to 0.10, and `VESPA-SS` remains first, with 0.807 against 0.773 of Severus (*s* = 8). The published call sets are within 0.02 of our default runs, except Sniffles2 on `HG008` and Severus on COLO829.

To further investigate these results, we stratified the ground truth calls by Variant Allele Frequency (VAF). For each true event, the VAF was estimated using svviz2 [28], and calls were partitioned into four groups: low VAF (< 0.10), low-to-moderate VAF (< 0.20), all calls with a reported VAF, and calls without a VAF estimate. Figure 3b summarizes the results for the low-to-moderate VAF group. Notably, on the `HG008` dataset, `VESPA-SS` achieves the highest F1 score across all tested values of *s*, exceeding the second-best caller by 9 to 21 percentage points. Interestingly, the increase in F1 score is larger when the evaluation is restricted to these more challenging variants. Overall, `VESPA-SS` detects the largest number of events while maintaining the highest precision. The only exception is *s* = 3, for which `VESPA-SS` correctly reports 17 events compared to 19 for Severus. However, Severus achieves substantially lower precision, reporting 72 false positives compared to 17 for `VESPA-SS`. Consequently, `VESPA-SS` remained the most precise caller even when using an extremely permissive support threshold. The results on the COLO829 dataset show a similar overall trend, although the differences between callers are less pronounced. Across all tested values of *s*, `VESPA-SS` consistently detects the highest number of true events, but lower support thresholds result in a decrease in precision. At low values of *s*, SAVANA achieves the highest F1 score, thanks to a more effective filtering of false positive calls. Increasing the minimum support threshold to *s* = 8, however, allows `VESPA-SS` to outperform all competing methods, achieving the highest F1 score (0.82), 12 percentage points higher than the best-performing state-of-the-art caller, SAVANA. In this configuration, `VESPA-SS` reports 20 true positives, 4 false negatives, and only 5 false positives. Overall, these results indicate that `VESPA-SS` outperforms state-of-the-art not only through increased sensitivity, but also through more effective filtering of false positive events.

Finally, we compared the events correctly reported by `VESPA-SS` with those reported by the other state-of-the-art approaches. For each method, we selected the configuration yielding the highest recall and compared the corresponding sets of true positive calls. The results of this analysis are shown in Figure 3c. On the `HG008` dataset, considering both clonal and subclonal regions, all callers show substantial agreement. Of the 173 true events, of which Minda evaluates 172, 106 are reported by all callers, whereas only 11 are missed by every method. Few events are reported by a single caller: one by SAVANA only, two by Severus, and four by Sniffles2 (whose performance, however, is substantially affected by its large number of false positives, 713). No event is uniquely reported by `VESPA-SS`. This result is not unexpected, as the `HG008` ground truth was generated from an ensemble that includes the four callers of our evaluation, whereas `VESPA-SS` did not contribute to its construction. Consequently, the evaluation is inherently biased toward these methods and their SV representations [5]. Nevertheless, `VESPA-SS` reports the highest total number of true events (153) while maintaining the lowest number of false positives (17). We note, however, that this analysis focuses exclusively on the highest-recall configuration of each caller and therefore does not account for the precision-recall trade-off. We therefore repeated this analysis considering the parameter configuration yielding the highest F1 score for each caller. As expected, the total number of events reported by the callers decreases. In this setting, seven events are reported by `VESPA-SS` only, whereas an additional seven events are reported exclusively by Severus and `VESPA-SS`. Together with its low false-positive rate, these results further highlight the optimal balance between precision and recall achieved by `VESPA-SS`. Similar results can be observed when considering the COLO829 dataset and the highest-recall configurations. In this case, all callers except for Sniffles2 were not involved in the creation of the ground truth, and `VESPA-SS` has been able to report two true events missed by all competing approaches, albeit with a somewhat higher number of false positives. The same trend can be observed when considering the configuration yielding the highest F1 score, where `VESPA-SS` continues to identify two events missed by every other caller.

Overall, these results suggest that the methodology implemented in `VESPA-SS` enables it to identify true somatic SVs missed by existing approaches while maintaining a competitive false positive rate, leading to consistently superior performance across datasets.

### 3.2 Advantages of somatic signal discovery via SFS and Draft Composite Germline Reference

In our second experimental evaluation, we analyzed the contribution of two key innovations introduced in `VESPA-SS`: the use of a Draft Composite Germline Reference and the alignment-free extraction of somatic signal using specific strings.

We first evaluated the effectiveness of our Draft Composite Germline Reference and assessed that `VESPA-SS` does not require a high-quality assembly of the patient’s germline genome. To this end, we considered the `HG008` dataset and compared the SV calling performance of `VESPA-SS` using three different reference sets for specific strings computation: (i) the reference sequence together with the germline contigs, (ii) our Draft Composite Germline Reference, and (iii) the reference sequence together with a manually curated, near telomere-to-telomere germline assembly [30]. In all three settings, `VESPA-SS` correctly called 152 of the 172 true events. Using the Draft Composite Germline Reference reduced the number of false positives from 9 (when indexing only the reference sequence and the germline contigs) to 6, yielding a precision comparable to that obtained with the curated germline assembly (5 false positives). We note that the Draft Composite Germline Reference extends the sequences indexed in (i) by adding only 7.7% of the normal reads. These results demonstrate that our Draft Composite Germline Reference is a cost-effective and practical alternative to current state-of-the-art approaches, achieving nearly identical somatic SV calling performance without requiring curated high-quality assemblies.

We then evaluated how effectively specific strings computed against the Draft Composite Germline Reference capture the somatic evidence. To this end, we intersected the regions of the genome covered by specific strings with the 321 true breakpoints contained in the `HG008` ground truth (with a 50bp tolerance on both sides of each breakpoint). Specific strings captured 309 of the 321 somatic breakpoints (96.3%) while covering 13.7 Mb of the reference genome, corresponding to 0.49% of the clonal and subclonal regions considered in the analysis. To further contextualize these results, we compared them against specific strings computed using either the reference genome alone or the curated germline assembly (as done previously). Notably, the curated assembly produced a larger number of specific strings without increasing the number of captured breakpoints. This suggests that the additional germline sequence provides a more detailed representation of the genome but contributes little additional somatic evidence for breakpoint detection. In contrast, when indexing the reference genome alone, the number of specific strings increased to 205.7 million whereas the number of captured breakpoints increases by only 9 (318 out of 321, 99.1%). This marginal gain is partly driven by chance and comes at the cost of a substantial reduction in specificity, since the specific strings computed this way cover 190.55 Mb of the reference genome, resulting in an almost 14× increase compared to the Draft Composite Germline Reference.

To further investigate these results, we stratified the 321 true breakpoints according to both the class of variant they come from and the stage of `VESPA-SS` they reached (Figure 4). The initial set of specific strings covered 309 breakpoints while missing 12 whereas the final set of candidate SVs considered by `VESPA-SS` missed 33 of them. We therefore analyzed the reasons underlying these results. Of these 33 breakpoints, 10 belong to subclonal duplications and complex rearrangements (“other breakend” in Figure 4) whose specific strings do not yield a candidate SV. An additional 4 breakpoints correspond to two subclonal deletions that are mistakenly corrected as sequencing errors by our approach due to their low support, that is 3 and 5 supporting tumor reads, respectively. The remaining 19 breakpoints belong to the “tandem-repeat length changes” class. In three cases, the corresponding specific strings are present but assigned to the incorrect repeat copy: they are placed from 76 to 235 bp away from the true breakpoint (thus exceeding the 50bp threshold used in this analysis). Due to their inherently repetitive nature, these variants represent the most challenging class for all the evaluated callers. Of the 32 breakpoints of this class, 17 belong to events found by no caller. Sniffles2 achieved the highest recall, recovering 14 of them whereas the other approaches recovered only 2 to 6.

**Figure 4.**
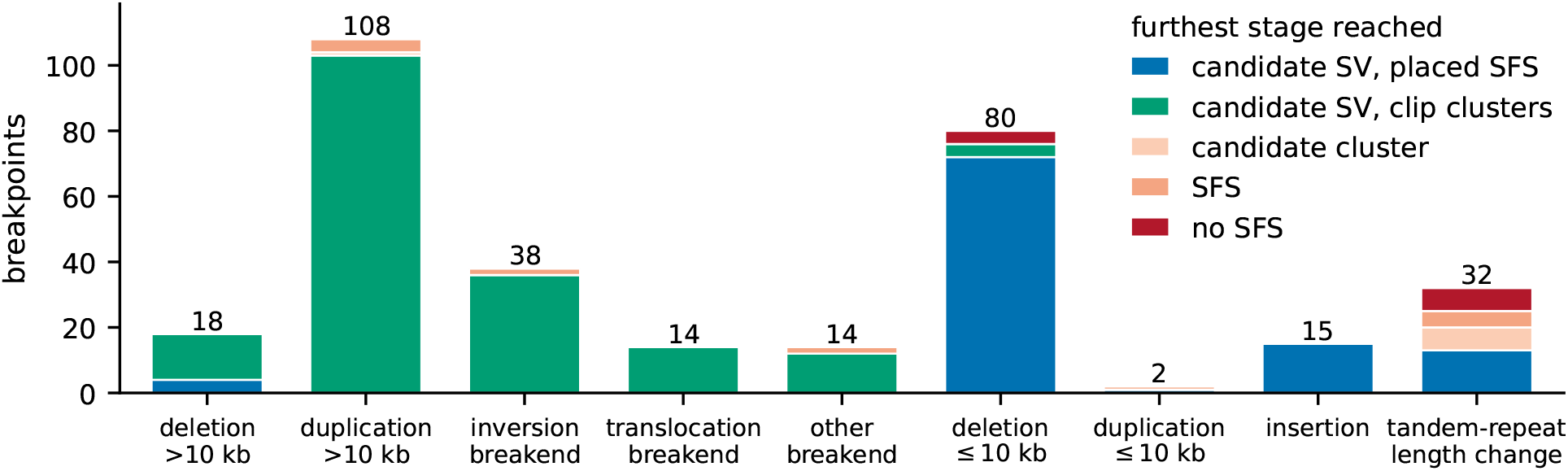
Breakpoints of the V0.5 ground truth on `HG008` (321, clonal and subclonal benchmark regions) by class of SV and by the furthest stage they reach with the full germline set. Candidate SVs are split by calling path; candidate cluster: a cluster supported by at least *s* = 5 reads; SFS: a placed SFS within 50 bp. Tandem-repeat length changes are the records the benchmark labels as copy-number changes of a tandem repeat; a breakend takes the class of the event the benchmark assigns it.

Figure 4 also highlights the complementary nature of the two calling paths implemented in `VESPA-SS`. Among the 288 breakpoints reached by a candidate SV, 63.5% are supported exclusively by clipped specific strings. This result demonstrates that the new calling strategy based on clipped specific strings implemented in `VESPA-SS` is essential for achieving competitive somatic SV calling accuracy. As expected, these breakpoints are associated with complex structural variants (such as long duplications, inversions, and translocations), which would be missed if relying solely on placed specific strings.

## 4 Discussion

We introduced `VESPA-SS`, a novel approach for calling somatic structural variants from paired tumor–normal long-read whole-genome sequencing data. By combining alignment-free extraction of somatic evidence against a Draft Composite Germline Reference with alignment-based calling procedures, `VESPA-SS` achieved state-of-the-art performance and outperformed existing alignment-based approaches. Across nearly all evaluated scenarios, `VESPA-SS` achieved the highest sensitivity while maintaining competitive precision. Notably, the largest gains were observed when calling structural variants with low-to-moderate variant allele frequencies, which represent one of the most challenging classes of somatic variants. `VESPA-SS` builds on our previous work on germline structural variants detection (SVDSS, [4]) but extends it substantially, resulting in a standalone somatic SV caller. Specifically, `VESPA-SS` introduces a somatic-aware error-correction strategy that preserves low-frequency somatic signals, novel calling procedures for simple and complex structural variants, and new probabilistic filtering procedures that improve its calling precision.

Among these contributions, a key innovation is the use of a Draft Composite Germline Reference to extract somatic evidence from the tumor sample while filtering germline signals without compromising sensitivity. This reasonably complete representation of the patient’s germline sequence enables `VESPA-SS` to achieve state-of-the-art accuracy without requiring the costly and demanding assembly of the germline genome of the patient, as adopted by several recent approaches [32, 19, 1]. As shown by our results, replacing a curated, near telomere-to-telomere germline assembly with the Draft Composite Germline Reference preserves the same number of true positive calls while increasing the number of false positives by only one event. These findings demonstrate the effectiveness of our approach and establish the Draft Composite Germline Reference as a practical and cost-effective alternative to current strategies. Future work will focus on further reducing its computational complexity while maintaining the ability to effectively separate somatic from germline signals. The main limitation of the current implementation lies in its ability to accurately resolve complex rearrangements. Each clip cluster contributes only a single junction (the one with highest support). Consequently, when multiple junctions share the same breakpoint, a common signature of complex rearrangements, only one of them is reported. This limitation explains why the missed events on the `COLO829` dataset are primarily this kind of complex events. Addressing this will be a major focus of future development. Future efforts will also be devoted to extend `VESPA-SS` to Oxford Nanopore Technologies sequencing data and perform extensive evaluation on low coverage paired tumor–normal samples. Based on the results presented here, we expect the advantages of `VESPA-SS` over current state-of-the-art approaches to become even more pronounced in these challenging settings.

